# Generation and characterization of a mouse model of conditional *Chd4* knockout in the endometrial epithelium

**DOI:** 10.1101/2025.06.05.658183

**Authors:** Shannon K. Harkins, Hilary J. Skalski, Abigail Z. Bennett, Laura Pavliscak, Amelia R. Arendt, Lauren Wood, Genna E. Moldovan, Ronald L. Chandler

## Abstract

Chromatin remodeling plays an integral part in endometrial homeostasis, having roles in the maintenance of cell identity, epithelial integrity, and prevention of endometrial disease. Chromodomain-helicase-DNA-binding protein 4 (*CHD4*) is a chromatin remodeling protein and member of the NuRD complex, which predominantly represses transcription. *CHD4* is mutated in endometrial carcinoma, with most mutations leading to loss of function. *CHD4* has been identified as a tumor suppressor and regulator of stemness in endometrial carcinoma cells, but little is known about the tissue-specific roles of *CHD4* in the endometrial epithelia *in vivo.* We generated a conditional *Chd4* floxed allele and combined it with *BAC-Sprr2f-Cre* to drive *Chd4* loss in the endometrial epithelium. Consistent with previous reports, *BAC-Sprr2f-Cre* expression is absent in the oviducts, ovaries, and kidneys, and it shows variegated expression within the endometrial epithelium. Loss of CHD4 was confirmed by immunohistochemistry, and stained cells were quantified to determine the percentage of endometrial epithelial cells with and without CHD4. Compared to the glandular epithelium, the extent of CHD4 loss was higher in the luminal epithelium and unaffected by age. Mice with conditional knockout of *Chd4* had normal endometrial histology. A 6-month breeding trial was performed to assess the functional effects of endometrial epithelial *Chd4* loss on fertility. No difference in litter size, mean number of pups per litter per dam, or pup weight was found between genotypes. These findings demonstrate that *Chd4* conditional loss using *BAC-Sprr2f-Cre* is not sufficient to alter the structure and function of the endometrial epithelium or drive tumorigenesis. As *CHD4* is frequently co-mutated with other cancer driver genes such as *TP53*, *PIK3CA*, and *PTEN*, future mouse modeling efforts emulating patient mutational profiles might provide insight into the role of *CHD4* in endometrial carcinoma.

## Introduction

The endometrium is a dynamic, hormone-responsive tissue that forms the inner uterine lining and consists of luminal epithelium, glandular epithelium, and stroma. The primary physiological function of the endometrium is to prepare for and maintain pregnancy(1). In response to ovarian steroid hormones, estrogen and progesterone, the endometrium undergoes cyclic proliferation and breakdown each menstrual cycle. Endometrial function, including the menstrual cycle and pregnancy, is tightly controlled by epigenetic regulation to maintain endometrial homeostasis(2–4). Loss of this epigenetic regulation can lead to endometrial pathologies, such as endometrial hyperplasia(5) and endometrial cancer(6).

Chromatin remodeling proteins play an integral role in the epigenetic regulation of the endometrium. Previous work from our lab identified an endometrial epithelial-specific role for SWI/SNF chromatin remodeling proteins, *ARID1A* and *BRG1,* in the regulation of epithelial cell identity and integrity, with loss of SWI/SNF leading to an epithelial-to-mesenchymal transition (EMT) phenotype(7, 8). Chromodomain-helicase-DNA-binding protein 4 (*CHD4*) is an ATP-dependent chromatin remodeling protein and integral subunit of the nucleosome remodeling and deacetylase (NuRD) complex, which regulates transcriptional repression, DNA damage response, and cell cycle progression (9–14). *CHD4* mutations are found in the benign and malignant endometrium(6, 15). CHD4 is mutated in 14.1% of endometrial carcinomas in the Cancer Genome Atlas Uterine Corpus Endometrial Carcinoma dataset(16). CHD4 is the predominant chromatin remodeling mutation in a rare and aggressive form of endometrial carcinoma known as uterine serous carcinoma, in which 17% of cases have CHD4 mutations(17, 18). CHD4 has been associated with the acquisition of a metastatic phenotype in several cancer types, including ovarian, colorectal, papillary thyroid, and breast cancers(19–22). We have previously identified a putative role for CHD4 in 12Z endometriotic epithelial cells in which CHD4 is recruited to H3.3-containing super-enhancers, where it co-represses genes associated with EMT, motility, adhesion, and epithelial cell identity in an ARID1A/SWI-SNF-dependent manner (23). Still, little is known about the tissue-specific functions of CHD4 within the normal endometrium *in vivo* or the extent to which CHD4 loss may contribute to endometrial disease. To this end, we generated a conditional *Chd4* floxed allele, which, when combined with *BAC-Sprr2f-Cre,* targets *Chd4* loss to the endometrial epithelia(24), the proposed cell of origin for endometrial carcinoma(25). We generated the first endometrial epithelial-specific conditional knockout of *Chd4* and characterized the structural and functional consequences of CHD4 loss.

## Materials and Methods

### Mice

All mice were maintained on a C57BL/6 background. The C57BL/6N-*Chd4^tm1a(EUCOMM)Wtsi^/BayMmucd* (knockout-first, conditional *Chd4* floxed-neo allele, *Chd4^fl-neo^*) allele was obtained from Mutant Mouse Resource and Research Centers at the University of California Davis (MMRRC #037690-UCD)(26). The *BAC-Sprr2f-Cre* (strain #: 037052), R26-Flp knock-in (*R26^Fki^*; strain #: 016226), and R26-mT/mG (*R26^mT/mG^*; strain #: 007676) alleles were purchased from The Jackson Laboratory (Bar Harbor, ME). Inheritance of the *BAC-Sprr2f-Cre*, the *R26^Fki^*, and the *R26^mT/mG^* alleles were confirmed by PCR using published methods (24, 27, 28). *BAC-Sprr2f-Cre-* negative floxed (fl/fl or fl/+) mice were used as control mice unless otherwise specified. All mouse experiments were performed under a protocol approved by the Michigan State University (MSU) Institutional Animal Care and Use Committee. Mice were housed at the MSU Grand Rapids Research Center Vivarium on a standard 12-hour light-dark cycle with ad libitum food and water.

### Generating the conditional *Chd4* floxed allele

The *Chd4^fl-neo^* allele configuration was generated through a targeted breeding scheme that induced sequential recombination by Flp1 and Cre-recombinase (Fig 1a.,c.-d.) The presence and configuration of the *Chd4^fl-neo^* allele were confirmed by PCR in accordance with MMRC-UCD genotyping protocol guidelines(Fig 1d.)(*29*). Table 1 summarizes the primer sequences, primer combinations, and PCR product sizes for each allele configuration of the *Chd4^fl-neo^* allele(*29*). Reactions highlighted in grey were not performed and are not included in Fig 1d. PCR reaction conditions consisted of initial denaturation at 94°C for 2 minutes, followed by 34 cycles at 94°C for 15 seconds, 58°C for 30 seconds, and 72°C for 1 minute, and ended with extension at 72°C for 5 minutes.

**Fig 1:**
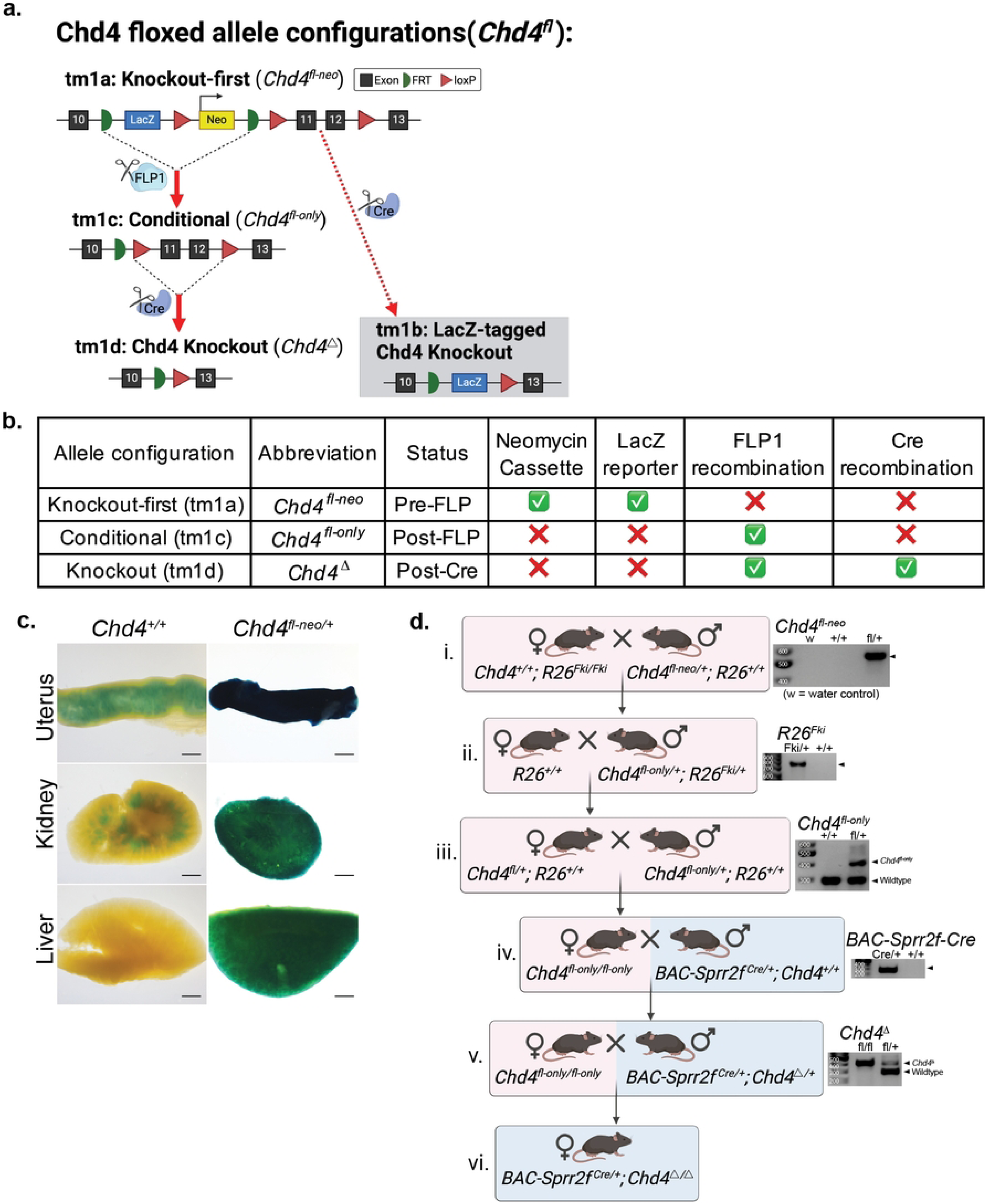
Generating the *Chd4* conditional knockout mouse. **a.** Schematic of the knockout-first, conditional Chd4 allele. The targeted mutation 1a (tm1a) allele configuration contains flippase recognition targets (FRT) flanking a *lacZ* reporter and a neomycin resistance cassette. LoxP sites flank the Neomycin cassette and critical exons 11 and 12. The tm1b allele configuration (gray box) is produced when Cre recombinase is introduced prior to FLP1 recombination (not generated in this study). The tm1c allele configuration is produced following FLP1 recombination. The tm1d allele configuration is only produced in cells expressing Cre recombinase. The use of *BAC-Sprr2f-Cre* drives conditional knockout of the Chd4 in the endometrial epithelium. **b.** Tabular summary of the tm1a, tm1c, and tm1d allele configurations for the *Chd4^fl-neo^* allele. **c.** X-gal staining was used to assess the functionality of the *Chd4^fl-neo^* in *Chd4^fl-neo^*^/+^ and control (Chd4^+/+^) mice. The *lacZ* reporter gene allows cells with the Chd4^fl-neo^ allele to produce β-galactosidase, which hydrolyses X-gal and produces a blue/green color. **d.** A selective breeding scheme was used to produce the conditional Chd4 knockout in the endometrial epithelium. Representative PCR genotyping results are displayed to the right of relevant steps containing each respective allele. i. *R26^Fki^* introduction induces FLP1 recombination in offspring that contain the *Chd4^fl-neo^* allele. ii. FLP1 recombination creates the *Chd4^fl-only^* allele. *R26^Fki^* is no longer needed and bred out by crossing with a wildtype C57BL/6 mouse. iii. A sire homozygous for the *Chd4^fl-only^* allele is crossed with a heterozygous dam to produce a dam homozygous for the *Chd4^fl-only^* ^allele^. iv. A sire containing the *BAC-Sprr2f-Cre* allele is crossed with a dam homozygous for the *Chd4^fl-only^* allele. v. BAC-Sprr2f-Cre drives Cre recombination in the sire, knocking out *Chd4* in Cre-expressing tissues and creating the *Chd4^Δ^* allele. The sire is crossed with a dam homozygous for the *Chd4^fl-only^* allele to create a dam homozygous for Chd4 loss in the endometrial epithelium (*Chd4^Δ^*).

**Table 1:**
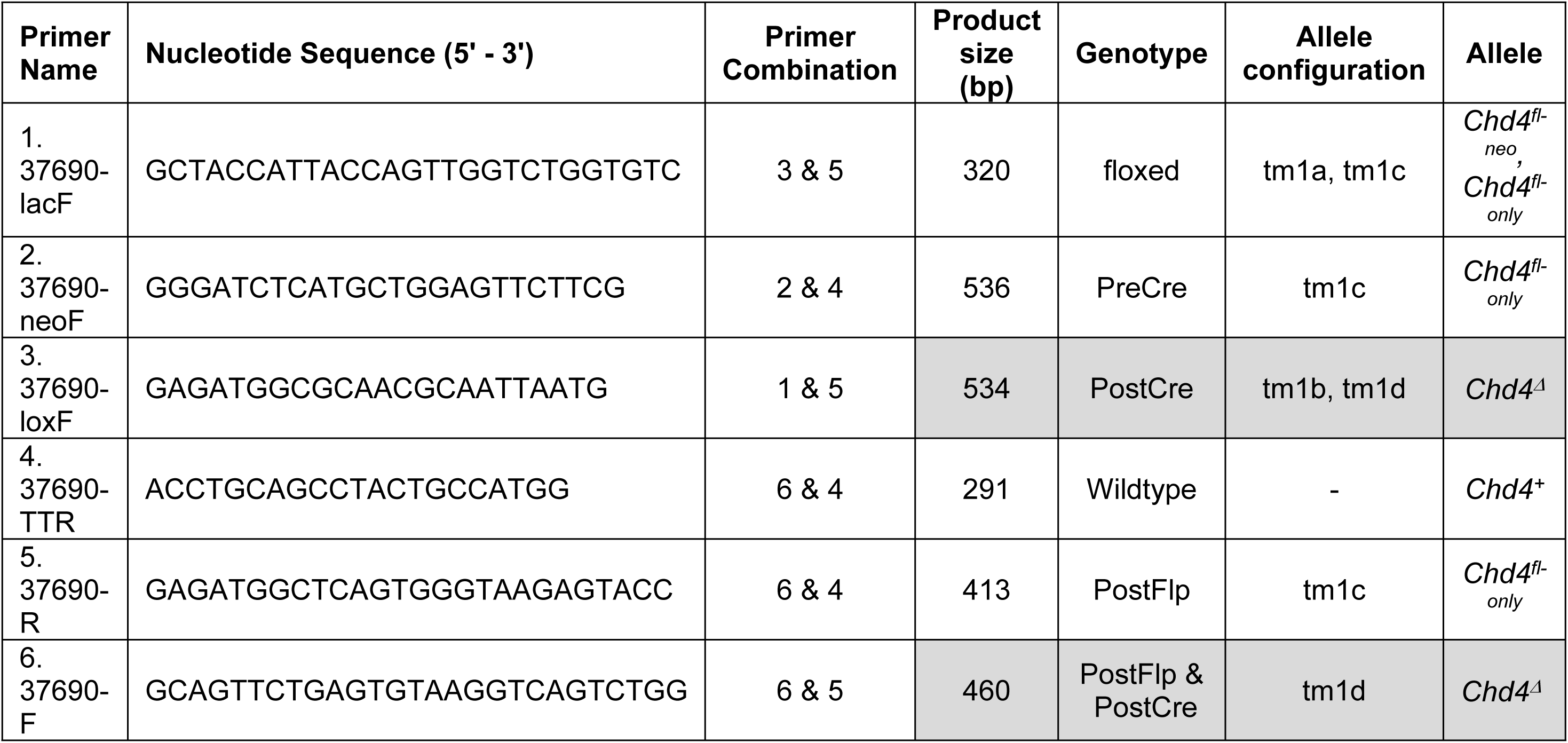
PCR Genotyping of the conditional *Chd4* floxed allele configurations:

### Estrous Cycle Staging

As described previously, vaginal lavage cytology was used to determine the murine estrous cycle stage was performed (30). Twelve-week-old pilot mice were not estrous cycle staged. Mice in the 26-week-old cohort were sacrificed in the diestrus phase of the estrous cycle.

### X-gal Staining

X-gal staining solution and wash buffer were prepared as described, except that the staining solution contained 20 mM Tris-HCL (pH 7.5)(31). All wash steps were repeated three times using wash buffer. Isolated organs were fixed in 4% paraformaldehyde (PFA) in phosphate-buffered saline (PBS) overnight at 4°C, washed, stained with X-gal staining solution overnight, washed, and placed in 4% PFA overnight for post-staining fixation. Samples were then washed and cleared through a graded glycerol series (50%, 70%, 90%, 100%) with consecutive overnight incubations.

### Whole Mount Tissue Imaging

Whole mount tissue samples, including X-gal stained tissues and *R26^mT/mG^* mice, were imaged using a Nikon SMZ1270i Stereo-Microscope and NIS-Elements Software (Nikon Instruments Inc., Tokyo, Japan).

### Tissue Processing and Histology

Upon necropsy, one uterine horn was fixed in 10% neutral buffered formalin in PBS for 72 hours. Samples were then washed twice with PBS and transferred to 70% ethanol. Samples were submitted to the Van Andel Research Institute Pathology & Biorepository Core for paraffin embedding, sectioning, and hematoxylin and eosin (H&E) staining. The other uterine horn was fixed in 4% paraformaldehyde (PFA) for at least 72 hours and stored at 4°C. Each uterine horn was washed twice with PBS, subjected to a sucrose gradient (24 hours in 15% sucrose in PBS followed by 24 hours in 30% sucrose in PBS), embedded in O.C.T Compound (Fisher Health Care), and cryosectioned as described(8).

### Quantification of CHD4 loss

For each mouse, a representative αCHD4 IHC image at 200x magnification was selected for quantification of endometrial epithelial CHD4 loss. The multipoint tool in Fiji/Image J was used to manually count endometrial epithelial cells with and without positive DAB staining for CHD4. Cell counts for the luminal and glandular epithelia were enumerated separately. The percentage of cells expressing CHD4 was calculated by dividing the number of positive cells by the total number of cells and multiplying by 100% for the luminal, glandular, and total (luminal + glandular) epithelia.

### Immunofluorescence (IF)

IF was performed as described(8). Slides were blocked using donkey blocking solution: 5% normal donkey serum (Jackson Immunoresearch Laboratories # 017-000-121), 1% IgG-free bovine serum albumin (Jackson Immunoresearch Laboratories # 001-000-161), and 0.05% Tween 20 in PBS). Cryosections were co-stained with chicken anti-GFP (Abcam #ab13970, 1:500) and rat anti-KRT8 (DSHB #TROMA1, 1:100). The following secondary antibodies were purchased from Jackson Immunoresearch Laboratories and used at a 1:250 dilution: donkey anti-chicken AF 488 (JAX #703-545-155) and donkey anti-Rat AF 647 (JAX #712-605-153). Autofluorescence reduction was performed using the TrueVIEW Auto-fluorescence Quenching Kit (Vector Laboratories #SP-8500). Slides were mounted and stained with DAPI using ProLong Gold Antifade Reagent with DAPI (Invitrogen). Slides were imaged using the Nikon C2 Plus Confocal microscope.

### Immunohistochemistry

Indirect immunohistochemistry (IHC) was performed as described(8, 30). Antigen retrieval was performed using 10mM sodium citrate buffer (pH 6).

Primary antibodies were incubated overnight at the following dilutions: 1:200 αCHD4 (Cell Signaling Technology, CST #12011), 1:250 αCleaved caspase 3 (CC3; CST # 9579), 1:400 αKi-67 (CST #12202), 1:100 αKeratin 8 (KRT8; Developmental Studies Hybridoma Bank #TROMA-I), 1:400 αE-Cadherin (CST# 3195), and 1:400 αGFP (CST#2956).

### Statistical Analysis

All statistical analyses were performed using GraphPad Prism 10. Scatter plots show the mean and standard deviation for each dataset. An unpaired, two-tailed t-test was used to assess the difference between genotypes unless otherwise specified. Welch’s Correction was used for samples with unequal variance. Significance thresholds: ns=not significant; p<0.05=*, p<0.01=**, p<0.001=***, p<0.0001=****.

## Results

### Creating the conditional *Chd4 flox-only* allele

The knockout-first, conditional *Chd4* floxed-neo allele (*Chd4^tm1a(EUCOMM)Wtsi^/Chd4^fl-neo^)* was obtained from the Mutant Mouse Resource and Research Center in the targeted mutation 1a(tm1a) configuration. The *Chd4^fl-neo^* allele contains a lacZ reporter and neomycin selection cassette flanked by flippase recognition target (FRT) sites and loxP sites flanking critical exons 11 and 12 (#037690-UCD)(26)(Fig 1a.). The presence of the *Chd4^fl-neo^* allele was confirmed by X-gal staining (Fig 1c.). The LacZ reporter present in the *Chd4^fl-neo^* allele hydrolyzes X-gal substrate, whose product produces a blue precipitate upon further oxidation (32). This color change was apparent in the uterus, kidney, and liver from the mouse containing the *Chd4^fl-neo^* allele (Fig 1c.). The uterus and the kidney had some background staining, which could result from endogenous β-galactosidase activity known to occur in these two organs(33).

A targeted breeding scheme was performed to achieve conditional knockout of *Chd4* exclusively in the endometrial epithelium driven by *BAC-Sprr2f-Cre* (JAX: #037052) (24). Representative agarose gels from PCR genotyping are displayed to the right of relevant steps containing each respective allele (Fig 1d.). The lacZ reporter and neomycin selection cassette present in the *Chd4^fl-neo^* allele are not needed *in vivo* and were removed to prevent off-target activity(34). The presence of the R26-Flp knock-in (*R26^Fki^*; JAX #016226) was confirmed by PCR (Fig 1d.ii.)(27). The first step toward FLP1 recombination was crossing a sire heterozygous for the *Chd4^fl-neo^* allele with a dam homozygous for *R26^Fki^* (Fig 1.d.i.). FLP1 recombination occurred in the resultant offspring that inherit both the *R26^Fki^ and Chd4^fl-neo^* alleles, producing the *Chd4^fl-only^* allele (Fig 1d.i.-ii.). To prevent future FLP1 recombination events, *Chd4^fl-only^; R26^Fki^* offspring were crossed with a wildtype C57BL/6 mouse to remove the *R26^Fki^* allele (Fig 1d.ii.).

The resultant offspring that were heterozygous for the *Chd4^fl-only^* allele were crossed to produce female *Chd4^fl-only^* homozygotes (Fig 1iii.-iv.). These female *Chd4^fl-only^* homozygotes were then crossed with a sire with a *Chd4* knockout driven by *BAC-Sprr2f-Cre* (Fig 1d.v.). The resultant female offspring with *BAC-Sprr2f-Cre* and homozygous for the conditional *Chd4^fl-only^* allele have CHD4 loss in the Cre-expressing cells of the endometrial epithelium (*BAC-Sprr2f^Cre/+^*; *Chd4*^Δ/Δ^) (Fig 1d.vi.).

### *BAC-Sprr2f-Cre* is expressed exclusively in the endometrial epithelium and is absent in the oviducts and ovaries

The *R26^mT/mG^* fluorescent Cre-reporter allele (JAX #: 007676) was used to characterize the activity of *BAC-Sprr2f-Cre* across the female reproductive tract in 12-week-old *BAC-Sprr2f^Cre/+^; R26^mT/mG^* and control (*BAC-Sprr2f^+/+^; R26^+/+^*) mice (Fig 2a.)(28). In mice with the *R26^mT/mG^* allele, all cells in the body express red fluorescent protein (RFP) except for the cells that express Cre recombinase, which express green fluorescent protein (GFP). Expression of the *BAC-Sprr2f-Cre* allele begins after the onset of puberty in a stochastic manner, with initial reports finding a 50% recombination in 6-week-old mice(24). In concordance with previously published results, *BAC-Sprr2f-Cre* was expressed exclusively in the endometrial epithelium, and expression was absent in the endometrial stroma, oviducts, and ovaries (Fig 2b.-c.)(24). The variegated expression characteristic of the *BAC-Sprr2f-Cre* allele is stronger in the luminal epithelium compared to the glandular epithelium by anti-GFP IHC in *BAC-Sprr2f^Cre/+^; R26^mT/mG^; Arid1a^fl/fl^* mice when compared to control mice (Fig 2c.).

**Figure 2:**
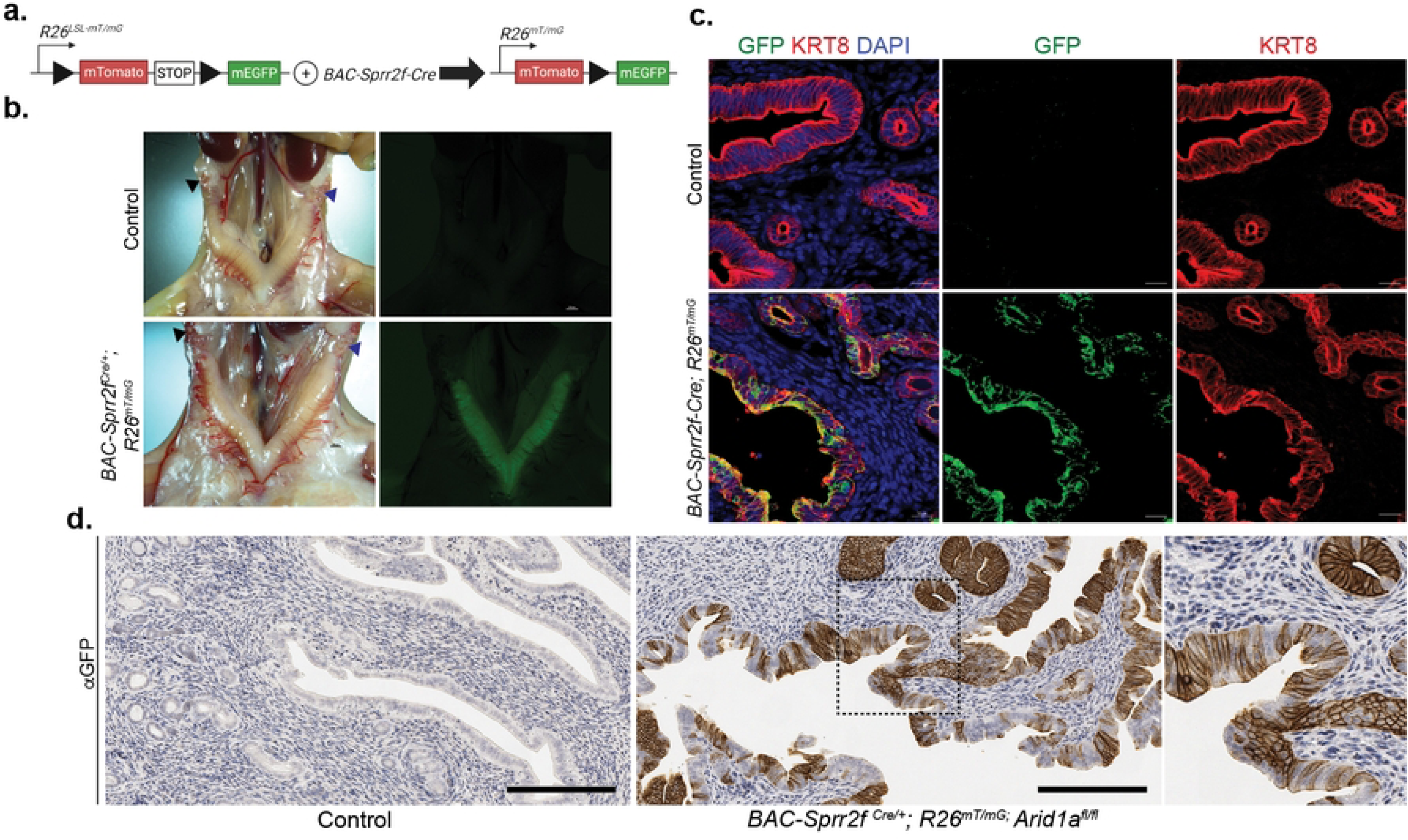
*BAC-Sprr2f-Cre* drives CRE activity in the endometrial epithelium. The R26-Lox-STOP-Lox (LSL)-mT/mG allele drives EGFP conditional expression in cells expressing *BAC-Sprr2f-Cre.* **b.** Photomicrograph of uteri from 12-week-old control (*BAC-Sprr2f^Cre/+^; R26^+/+^*) and *BAC-Sprr2f-Cre; R26^mT/mG^* and *BAC-Sprr2f^Cre/+^; R26^mT/mG^* mice. Cre-positive cells within the endometrial epithelium fluoresce green in mT/mG mutant mice. Cre expression is absent in the oviducts and ovaries. Scale bar=100 µm. **c.** Immunofluorescence staining for epithelial marker αkeratin 8 (KRT8), αGFP, and DAPI nuclear stain was performed on cryosections of the uteri in **b.** In the three-color merged image, co-expression is shown as yellow. The *BAC-Sprr2f-Cre* allele expression drives *R26^mT/mG^* expression exclusively in the endometrial epithelium and is absent in the stroma. Maximum intensity projection images at 400x magnification; scale bar=10 µm. **d.** Anti-GFP immunohistochemistry of formalin fixed paraffin embedded uteri from 15-week-old control (*BAC-Sprr2f ^+/+^; R26^+/+^; Arid1a^fl/fl^*) and *BAC-Sprr2f ^Cre/+^; R26^mT/mG^; Arid1a^fl/fl^* shows variegated patterns of brown DAB staining in the luminal and glandular epithelia. Images were taken at 200x magnification with 400x magnification in the inset image (far right); Scale bar=200 µm.

### *BAC-Sprr2f-Cre* drives variegated loss of *Chd4* exclusively in the endometrial epithelium

We have created a novel mouse model of endometrial epithelial-specific *Chd4* loss driven by *BAC-Sprr2f-Cre*. For the initial pilot cohort, three *Chd4* conditional knockout (cKO, *BAC-Sprr2f^Cre/+^; Chd4^/fl^*) mice and three Cre-negative control (*BAC-Sprr2f^+/+^; Chd4^/fl^* or *Chd4^/+^)* mice were sacrificed at 12 weeks of age to confirm loss of *Chd4* in endometrial epithelium and to assess endometrial histology. Pilot mice were not estrous cycle staged. Control mice showed strong nuclear expression of CHD4 in the luminal and glandular epithelia (Fig 3a.). The *Chd4* cKO mice show variegated expression of CHD4 in the luminal and glandular epithelia. The extent of CHD4 loss was determined from αCHD4 IHC and was quantified using Fiji/ImageJ to manually count DAB-positive and DAB-negative cell populations within the luminal and glandular epithelia. The percentage of cells expressing CHD4 was calculated as the number of DAB-positive cells divided by the total number of cells multiplied by 100% and was quantified for the luminal, glandular, and total (luminal + glandular) epithelia (Fig 3b.-3h.). The percent knockout is then 100% minus the percentage of cells expressing CHD4. Table 2 summarizes these values. The luminal epithelium had roughly 2-fold more CHD4 loss than the glandular epithelium in all three *Chd4* cKO mice (Fig 3e. KO1-3). The extent of CHD4 loss in the 12-week-old mice was consistent with what has been reported in the literature in terms of *BAC-Sprr2f-Cre* expression(24).

**Figure 3:**
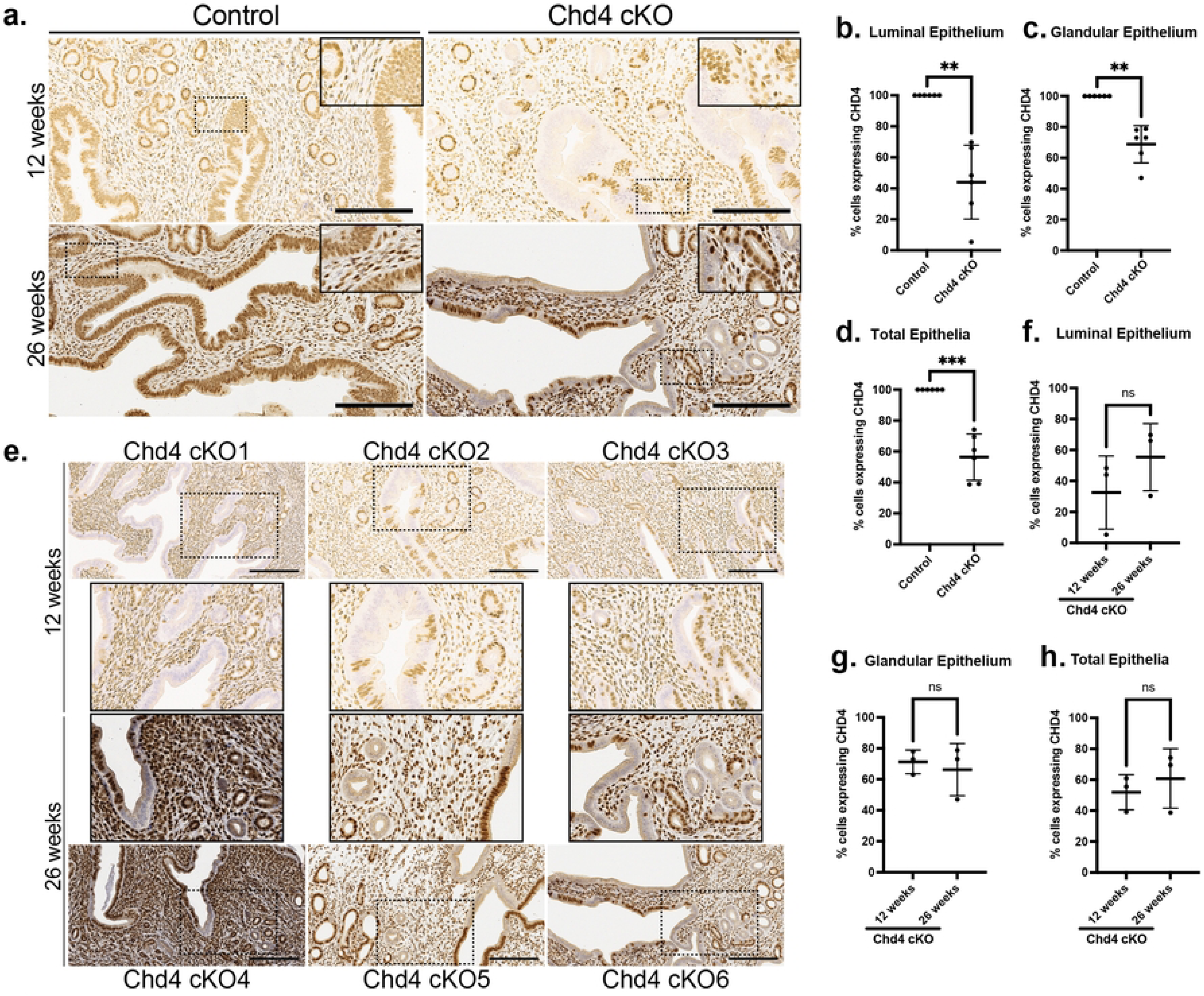
*BAC-Sprr2f-Cre* drives variegated loss of CHD4 in the endometrial epithelia. **a.** CHD4 loss was assessed by anti-CHD4 immunohistochemistry (IHC). Representative images of the endometria of 12 and 26-week-old mice Chd4 KO and control mice. **b.-h.** The percentage of CHD4-expressing cells within the endometrial epithelium (DAB-positive by anti-CHD4 IHC) as counted in Fiji/ImageJ. The percentage of CHD4-expressing cells was calculated as: the number of positive cells/total number of cells*100% and was calculated for the luminal, glandular, and total epithelial cells. **b.-d.** Comparison of the percentage of CHD4 expressing cells between *Chd4* cKO (n=6) and control (n=6) mice. **e.** Representative images of αCHD4 IHC showing the variation in Chd4 cKO in the 12-week old (cKO1-3) and 26-week-old (cKO4-6) mice. Images were taken at 200x magnification. The dashed box in the 200x image is enlarged to 400x magnification in the adjacent inset. **f.-g.** Comparison of the percentage of CHD4 expression between 12 and 26-week-old Chd4 cKO (n=6) and control (n=6) mice. The statistic used was a two-tailed, unpaired t test. Welch’s Correction was only used for unequal variance. Significance thresholds: ns=not significant; p<0.05=*, p<0.01=**, p<0.001=***, p<0.0001=****.

**Table 2:** Mean percentage of endometrial epithelial cells with and without CHD4 protein expression.

| Epithelium | Genotype | % cells expressing CHD4 | % cells with CHD4 Loss | p value |
| --- | --- | --- | --- | --- |
| Luminal | Control | 100 | 0 | 0.0022 |
|  | *Chd4* cKO | 44 | 56.1 |  |
| Glandular | Control | 100 | 0 | 0.0014 |
|  | *Chd4* cKO | 68.8 | 31.2 |  |
| Total | Control | 100 | 0 | 0.0008 |
|  | *Chd4* cKO | 56.4 | 43.6 |  |

Knowing that *BAC-Sprr2f-*Cre allele expression is estrogen-dependent, we next evaluated the extent to which CHD4 loss progressed during the aging process and cumulative exposure to endogenous estrogen(24). The histology and extent of CHD4 loss were assessed in a cohort of 26-week-old (6 months old) diestrus staged *Chd4* cKO (n=3) and control (n=3) mice. As with the 12-week-old pilot cohort, the 26-week-old *Chd4* cKO mice had a stronger degree of CHD4 loss in the luminal epithelium compared to the glandular epithelium (Fig 3b. KO4-6). No difference in the percentage of CHD4-expressing cells was detected between the 12-week-old and 26-week-old mice (Fig 3f.-h.). All six *Chd4* cKO and control mice showed a significant decrease in CHD4-expressing cells in the luminal, glandular, and total (average of the two) epithelia (Fig 3b.-h.). The luminal epithelium showed fewer CHD4-expressing cells than the glandular epithelium in all six *Chd4* cKO mice (Fig 3e.-f.). We did not note any apparent histologic differences between *Chd4* cKO and control mice at 12 weeks or 26 weeks of age (Fig 4a.). There was no difference in the expression of epithelial (E-cadherin), proliferative (Ki-67), or apoptotic (CC3) markers between genotypes (Fig 4b).

**Figure 4:**
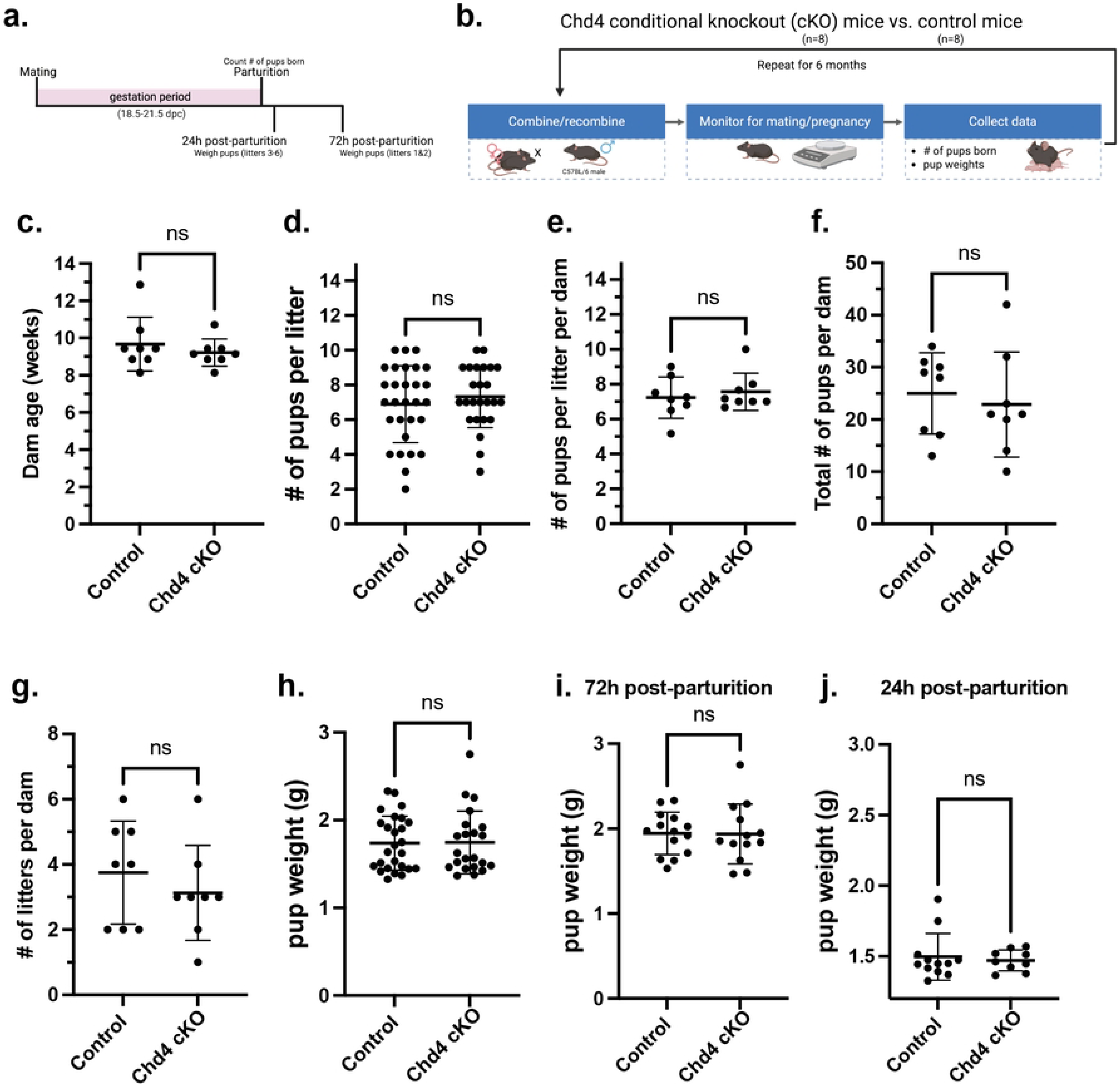
Chd4 cKO mice have normal histopathology. **a.** Representative images of hematoxylin and eosin stained endometria from 12 and 26-week-old mice. Representative images of IHC from 12-week-old mice assessing the expression of epithelial (E-cadherin), proliferative (Ki-67), and apoptotic (Cleaved Caspase 3/CC3) markers between genotypes. For images taken at 40x and 200x magnification, the scale bar= 600 and 200 µm, respectively.

### *Chd4* cKO mice are fertile and have no difference in fertility when compared to controls

The functional effects of Chd4 loss on endometrial function were assessed by a 6-month breeding trial in which *Chd4* cKO (n=8) and control (n=8) dams were serially bred to a C57BL/6 sire (Fig 5a.-b.). Over the course of the breeding trial, all *Chd4* cKO and control dams successfully mated and gave birth to live pups. Dam age ranged from 8 to 14 weeks of age at the start of the trial, and there was no significant difference in the dam age between genotypes (Fig 5c.). There was no difference in the number of pups per litter (Fig 5d.), the mean number of pups per litter per dam(Fig 5e.), the total number of pups per dam (Fig 5f.), the number of litters per dam(Fig 5g.), or mean pup weight between genotypes (Fig 5h.-j.). Figure 5a summarizes the two time points at which pup weight was collected: 72 hours post-parturition for litters 1-2 and 24 hours post-parturition for litters 3-6. There was no difference in pup weight at either timepoint (Fig 5i.-j.) or when combined (Fig 5h.). Table 3 summarizes these results. Statistic was a two-tailed, unpaired t test.

**Figure 5:**
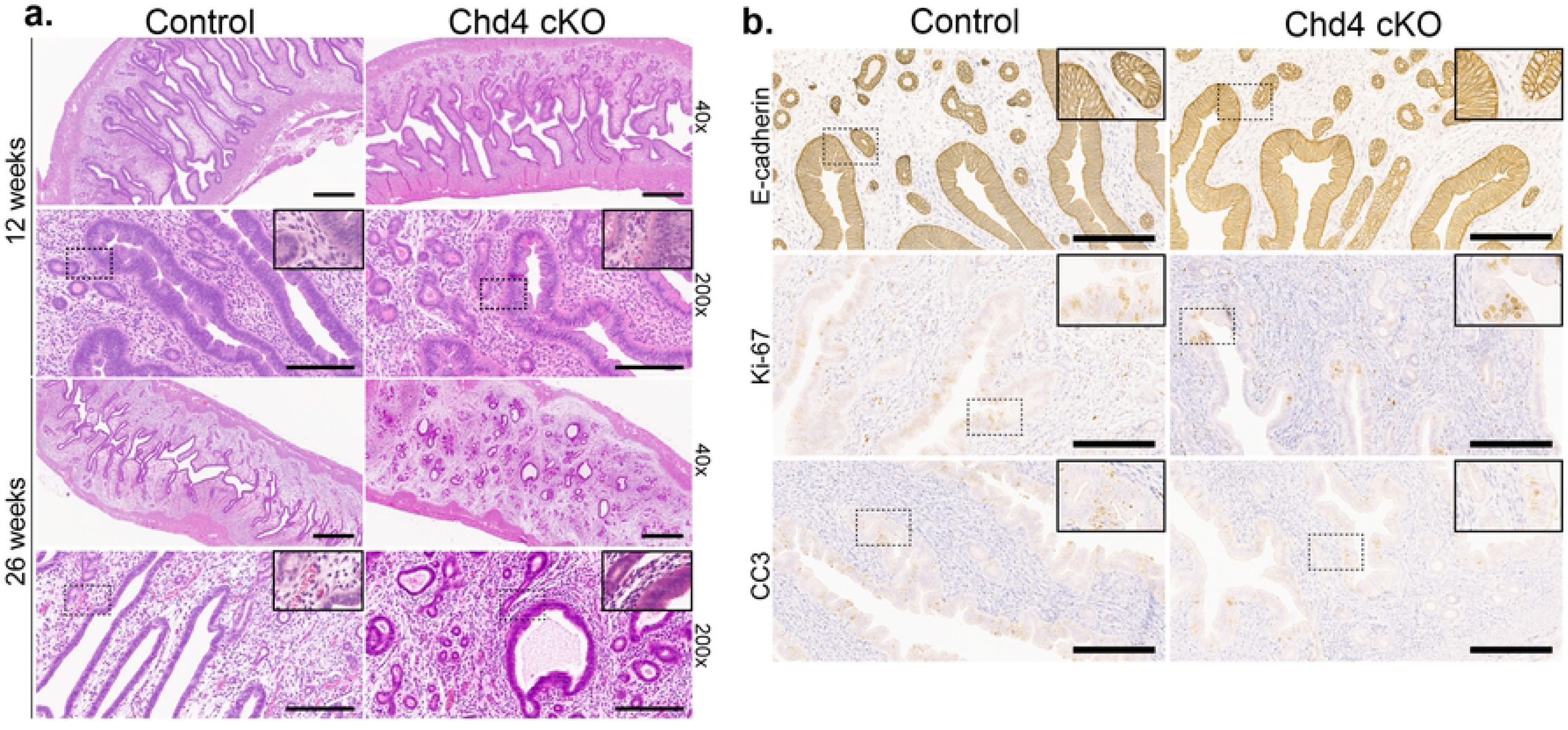
*Chd4* conditional knockout mice show no difference in fertility compared to control mice. **a.** Representative timeline of mouse pregnancy. Observed gestation periods ranged from 18.5 to 21.5 days. **b.** Visual summary of experimental design for the Chd4 breeding trial. **c.** There was no difference in dam age at the start of the breeding trial. **d.** *Chd4* KO dams have significantly fewer pups born per litter on average than control dams. Each datapoint represents one litter. **e.** The average number of pups per litter per *Chd4* KO dam was significantly less than control dams. Each datapoint represents one dam. **f.-g.** There was no difference in the total number of pups per dam **(f.)** or the number of litters per dam between genotypes (**g.). h.** There was no difference in pup weight between genotypes. As shown in a., litters 1 and 2 were weighed 72 hours post-parturition (**i.**), whereas litters 3 to 6 were weighed 24 hours post-parturition(**j.**). The statistic used was a two-tailed, unpaired t-test. Welch’s Correction was only used for unequal variance. Significance thresholds: ns=not significant.

**Table 3:**
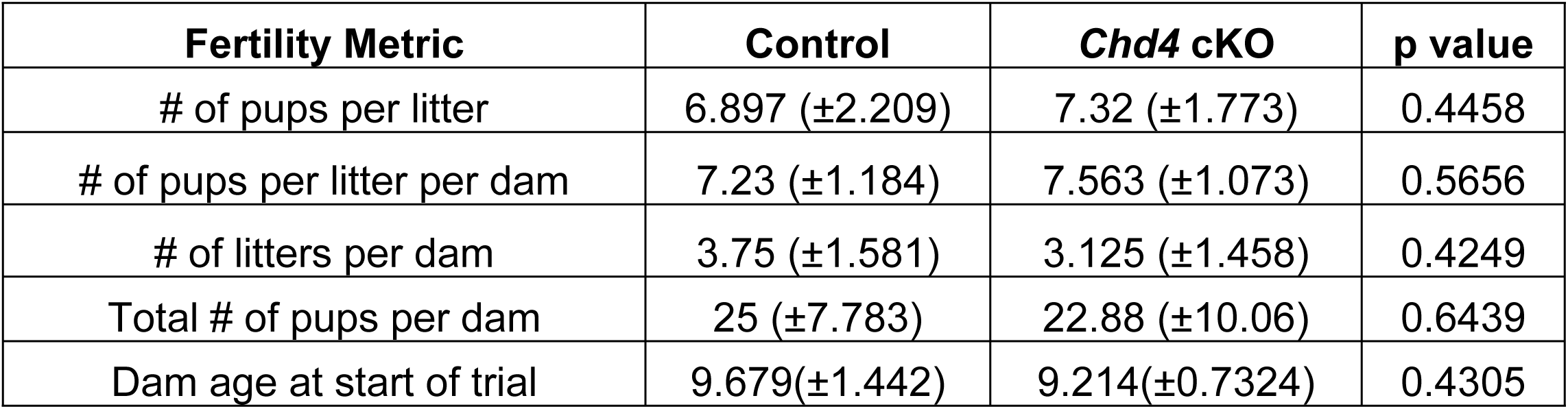
Breeding Trial Summary.

| Fertility Metric | Control | *Chd4* cKO | p value |
| --- | --- | --- | --- |
| # of pups per litter | 6.897 (±2.209) | 7.32 (±1.773) | 0.4458 |
| # of pups per litter per dam | 7.23 (±1.184) | 7.563 (±1.073) | 0.5656 |
| # of litters per dam | 3.75 (±1.581) | 3.125 (±1.458) | 0.4249 |
| Total # of pups per dam | 25 (±7.783) | 22.88 (±10.06) | 0.6439 |
| Dam age at start of trial | 9.679(±1.442) | 9.214(±0.7324) | 0.4305 |

## Discussion

Somatic mutations in CHD4 are found in the endometria of healthy women and those with endometrial cancer(6, 15). Knowing that endometrial carcinomas and other pathologies of the endometrium, including endometrial hyperplasia(5) and endometriosis(35), are thought to arise from the endometrial epithelium, we created an endometrial epithelial-specific mouse model of conditional *Chd4* loss. Through targeted breeding, FLP1 recombination excised the FRT site-flanked *lacZ* reporter and Neomycin cassette, creating the *Chd4^fl-only^* allele. At the onset of puberty, *BAC-Sprr2f-Cre* induced Cre recombination of the *Chd4^fl-only^* allele, leading to conditional loss of *Chd4* in the Cre-expressing cells of the endometrial epithelium. Mice with conditional *Chd4* loss exhibited variegated expression pattern characteristic of *BAC-Sprr2f-Cre* activity reported in the literature(24, 36). Although variegated, *Chd4* loss was more extensive in the luminal epithelium than in the glandular epithelium and did not change with age. *Chd4* cKO mice did not show any gross anatomical or histological differences when compared to control mice. Endometrial function was assessed by a 6-month breeding trial, which showed no difference in fertility between genotypes. Collectively, these results demonstrate that variegated *Chd4* loss does not alter endometrial structure or function in 6-month-old mice. Is this lack of phenotype due to the incomplete knockout caused by variegation, or is *Chd4* dispensable for endometrial epithelial function?

In mice, germline knockout of *Chd4* knockout (Chd4^-/-.^) is embryonic lethal due to implantation failure(37). In contrast, *Chd4^+/-^* (heterozygous) mice survive into adulthood but show altered growth, as well as altered neurological, cardiovascular, and reproductive development(38), similar to what is observed in Sifrim-Hitz-Weiss syndrome in humans(39, 40). Importantly, mice heterozygous for CHD4 loss can still reproduce, which may explain why the *Chd4* cKO mice with 56.4% remaining CHD4 expression in endometrial epithelia exhibited normal fertility. While CHD4 expression is haploinsufficient during development, the consequences of post-developmental *CHD4* loss appear to be tissue-specific. In malignancy, *CHD4* has context-dependent functions as a tumor suppressor and an oncogene. *CHD4* overexpression is associated with oncogenic activity, poor prognosis, and increased risk of colorectal cancer and ovarian cancer metastases(20, 22). In endometrial carcinoma, *CHD4* has been reported to have both tumor suppressive(41) and oncogenic activity(42), supporting the notion that *CHD4* gene dosage may be key to understanding the context-dependent CHD4 activity.

*BAC-Sprr2f-Cre* was selected due to its reported specificity to the endometrial epithelium and absence in the endometrial stroma, myometrium, ovaries, oviducts, and kidneys(24). To elucidate the extent to which variegated CHD4 loss contributed to the observed absence of a phenotype, an alternative endometrial epithelial-specific Cre driver, such as Lactoferrin i-Cre (*Ltf-iCre*), could be used to further evaluate the extent to which endometrial epithelial CHD4 is required for endometrial epithelial structure and function(43). However, lactoferrin is expressed in other tissues, including mammary glands and neutrophils, the off-target effects of which may alter endometrial functions such as fertility(43, 44).

CHD4 is a member of the CHD subfamily II proteins together with CHD3 and CHD5. The three proteins share high levels of amino acid sequence similarity, are members of distinct NuRD complexes, and play a role in the DNA damage response(45). CHD5 expression is confined to the brain and male reproductive tract, while CHD4 and CHD3 are ubiquitously expressed(46). In some contexts, distinct and redundant functions have been reported for CHD3 and CHD4(46). In other contexts, such as in murine cortical development, distinct NuRD configurations with CHD3, CHD4, and CHD5 have different functions(47). It remains possible other CHD proteins are compensating for CHD4 loss?

The majority of CHD4 mutations in endometrial carcinoma are missense mutations and are thought to lead to reduced or loss of CHD4 function(16, 41, 48). By targeting *Chd4* loss to the endometrial epithelium, we took a functional approach both to model the purported effects of these mutations and understand the tissue-specific consequences of CHD4 in this cell type. Phenotypic differences between the malignant behavior of tumors with TP53 mutations and TP53 loss have been well characterized, with certain mutations having gain-of-function effects that increase tumorigenicity(49). Accordingly, it is conceivable the consequences of a CHD4 mutation in tumors, even a loss-of-function mutation, may not be entirely congruent with the effects of a complete knockout, as modeled here.

Endometrial carcinoma most commonly arises in peri– and post-menopausal women(16). The mice used in this study were still cycling and therefore, may not have represented the atrophic endometrial epithelium common in post-menopausal women. Further, CHD4 is always co-mutated with other cancer driver genes and is frequently co-mutated with *PTEN*, *PIK3CA*, *PIK3R1*, *ARID1A*, *TP53*, and *KRAS*(16). Many chromatin remodeler mutations are necessary but not sufficient to drive endometrial tumorigenesis, and perhaps CHD4 needs an additional genetic assault or activating mutation for tumorigenesis to occur(8, 50, 51).

## Acknowledgments

We thank the Van Andel Research Institute Histology and Pathology Core for their histology and pathology services. We also thank Michigan State University’s Grand Rapids Research Center Vivarium for their assistance with our mouse care. Research reported in this publication was supported in part by grants from the Eunice Kennedy Shriver National Institute of Child Health & Human Development of the National Institute of Health under Award Numbers R01HD103617 and T32HD087166, MSU AgBio Research, and Michigan State University. The content is solely the responsibility of the authors and does not necessarily represent the official views of the National Institutes of Health.

## Contributions

Conceptualization: S.K.H. and R.L.C.; Methodology: S.K.H., R.L.C, and G.E.M..; Investigation: S.K.H., H.J.S., A.Z.B., L.P., A.A., L.W., G.E.M., and R.L.C.; Formal analysis: S.K.H., H.J.S., G.E.M.; Writing—original draft preparation: S.K.H. Writing— review and editing: H.J.S., A.Z.B., L.P., A.A., L.W., G.E.M., and R.L.C.; Funding acquisition and supervision: R.L.C.

## References

1. Critchley HOD, Maybin JA, Armstrong GM, Williams ARW. Physiology of the Endometrium and Regulation of Menstruation. Physiol Rev. 2020;100(3):1149–79.

2. Munro SK, Farquhar CM, Mitchell MD, Ponnampalam AP. Epigenetic regulation of endometrium during the menstrual cycle. Mol Hum Reprod. 2010;16(5):297–310.

3. Vrljicak P, Lucas ES, Tryfonos M, Muter J, Ott S, Brosens JJ. Dynamic chromatin remodeling in cycling human endometrium at single-cell level. Cell Rep. 2023;42(12):113525.

4. Retis-Resendiz AM, Gonzalez-Garcia IN, Leon-Juarez M, Camacho-Arroyo I, Cerbon M, Vazquez-Martinez ER. The role of epigenetic mechanisms in the regulation of gene expression in the cyclical endometrium. Clin Epigenetics. 2021;13(1):116.

5. Russo M, Newell JM, Budurlean L, Houser KR, Sheldon K, Kesterson J, et al. Mutational profile of endometrial hyperplasia and risk of progression to endometrioid adenocarcinoma. Cancer. 2020;126(12):2775–83.

6. Momeni-Boroujeni A, Vanderbilt C, Yousefi E, Abu-Rustum NR, Aghajanian C, Soslow RA, et al. Landscape of chromatin remodeling gene alterations in endometrial carcinoma. Gynecol Oncol. 2023;172:54–64.

7. Reske JJ, Wilson MR, Holladay J, Wegener M, Adams M, Chandler RL. SWI/SNF inactivation in the endometrial epithelium leads to loss of epithelial integrity. Hum Mol Genet. 2020;29(20):3412–30.

8. Wilson MR, Reske JJ, Holladay J, Wilber GE, Rhodes M, Koeman J, et al. ARID1A and PI3-kinase pathway mutations in the endometrium drive epithelial transdifferentiation and collective invasion. Nat Commun. 2019;10(1):3554.

9. Musselman CA, Ramirez J, Sims JK, Mansfield RE, Oliver SS, Denu JM, et al. Bivalent recognition of nucleosomes by the tandem PHD fingers of the CHD4 ATPase is required for CHD4-mediated repression. Proc Natl Acad Sci U S A. 2012;109(3):787–92.

10. Tong JK, Hassig CA, Schnitzler GR, Kingston RE, Schreiber SL. Chromatin deacetylation by an ATP-dependent nucleosome remodelling complex. Nature. 1998;395(6705):917–21.

11. Xue Y, Wong J, Moreno GT, Young MK, Cote J, Wang W. NURD, a novel complex with both ATP-dependent chromatin-remodeling and histone deacetylase activities. Mol Cell. 1998;2(6):851–61.

12. Zhang Y, LeRoy G, Seelig HP, Lane WS, Reinberg D. The dermatomyositis-specific autoantigen Mi2 is a component of a complex containing histone deacetylase and nucleosome remodeling activities. Cell. 1998;95(2):279–89.

13. Basta J, Rauchman M. The nucleosome remodeling and deacetylase complex in development and disease. Transl Res. 2015;165(1):36–47.

14. O’Shaughnessy A, Hendrich B. CHD4 in the DNA-damage response and cell cycle progression: not so NuRDy now. Biochem Soc Trans. 2013;41(3):777–82.

15. Kyo S, Sato S, Nakayama K. Cancer-associated mutations in normal human endometrium: Surprise or expected? Cancer Sci. 2020;111(10):3458–67.

16. Cancer Genome Atlas Research N, Kandoth C, Schultz N, Cherniack AD, Akbani R, Liu Y, et al. Integrated genomic characterization of endometrial carcinoma. Nature. 2013;497(7447):67–73.

17. Le Gallo M, O’Hara AJ, Rudd ML, Urick ME, Hansen NF, O’Neil NJ, et al. Exome sequencing of serous endometrial tumors identifies recurrent somatic mutations in chromatin-remodeling and ubiquitin ligase complex genes. Nat Genet. 2012;44(12):1310–5.

18. Zhao S, Choi M, Overton JD, Bellone S, Roque DM, Cocco E, et al. Landscape of somatic single-nucleotide and copy-number mutations in uterine serous carcinoma. Proc Natl Acad Sci U S A. 2013;110(8):2916–21.

19. Luo CW, Wu CC, Chang SJ, Chang TM, Chen TY, Chai CY, et al. CHD4-mediated loss of E-cadherin determines metastatic ability in triple-negative breast cancer cells. Exp Cell Res. 2018;363(1):65–72.

20. Wang J, Zhong F, Li J, Yue H, Li W, Lu X. The epigenetic factor CHD4 contributes to metastasis by regulating the EZH2/beta-catenin axis and acts as a therapeutic target in ovarian cancer. J Transl Med. 2023;21(1):38.

21. Pratheeshkumar P, Siraj AK, Divya SP, Parvathareddy SK, Alobaisi K, Al-Sobhi SS, et al. CHD4 Predicts Aggressiveness in PTC Patients and Promotes Cancer Stemness and EMT in PTC Cells. Int J Mol Sci. 2021;22(2).

22. Xia L, Huang W, Bellani M, Seidman MM, Wu K, Fan D, et al. CHD4 Has Oncogenic Functions in Initiating and Maintaining Epigenetic Suppression of Multiple Tumor Suppressor Genes. Cancer Cell. 2017;31(5):653–68 e7.

23. Reske JJ, Wilson MR, Armistead B, Harkins S, Perez C, Hrit J, et al. ARID1A-dependent maintenance of H3.3 is required for repressive CHD4-ZMYND8 chromatin interactions at super-enhancers. BMC Biol. 2022;20(1):209.

24. Cuevas IC, Sahoo SS, Kumar A, Zhang H, Westcott J, Aguilar M, et al. Fbxw7 is a driver of uterine carcinosarcoma by promoting epithelial-mesenchymal transition. Proc Natl Acad Sci U S A. 2019;116(51):25880–90.

25. Makker V, MacKay H, Ray-Coquard I, Levine DA, Westin SN, Aoki D, et al. Endometrial cancer. Nat Rev Dis Primers. 2021;7(1):88.

26. Skarnes WC, Rosen B, West AP, Koutsourakis M, Bushell W, Iyer V, et al. A conditional knockout resource for the genome-wide study of mouse gene function. Nature. 2011;474(7351):337–42.

27. Farley FW, Soriano P, Steffen LS, Dymecki SM. Widespread recombinase expression using FLPeR (flipper) mice. Genesis. 2000;28(3-4):106–10.

28. Muzumdar MD, Tasic B, Miyamichi K, Li L, Luo L. A global double-fluorescent Cre reporter mouse. Genesis. 2007;45(9):593–605.

29. (MMRRC) MMRRC. Genotyping protocol for Stock No. 037690: University of California, Davis; n.d. [Available from: https://mmrrc.ucdavis.edu/protocols/037690Geno_Protocol.pdf.

30. Skalski HJ, Arendt AR, Harkins SK, MacLachlan M, Corbett CJM, Goy RW, et al. Key Considerations for Studying the Effects of High-Fat Diet on the Nulligravid Mouse Endometrium. J Endocr Soc. 2024;8(7):bvae104.

31. DiLeone RJ, Russell LB, Kingsley DM. An extensive 3’ regulatory region controls expression of Bmp5 in specific anatomical structures of the mouse embryo. Genetics. 1998;148(1):401–8.

32. Blanco MJ, Learte AIR, Marchena MA, Munoz-Saez E, Cid MA, Rodriguez-Martin I, et al. Tracing Gene Expression Through Detection of beta-galactosidase Activity in Whole Mouse Embryos. J Vis Exp. 2018(136).

33. Merkwitz C, Blaschuk O, Schulz A, Ricken AM. Comments on Methods to Suppress Endogenous beta-Galactosidase Activity in Mouse Tissues Expressing the LacZ Reporter Gene. J Histochem Cytochem. 2016;64(10):579–86.

34. Pham CT, MacIvor DM, Hug BA, Heusel JW, Ley TJ. Long-range disruption of gene expression by a selectable marker cassette. Proc Natl Acad Sci U S A. 1996;93(23):13090–5.

35. Marquardt RM, Tran DN, Lessey BA, Rahman MS, Jeong JW. Epigenetic Dysregulation in Endometriosis: Implications for Pathophysiology and Therapeutics. Endocr Rev. 2023;44(6):1074–95.

36. Sahoo SS, Ramanand SG, Gao Y, Abbas A, Kumar A, Cuevas IC, et al. FOXA2 suppresses endometrial carcinogenesis and epithelial-mesenchymal transition by regulating enhancer activity. J Clin Invest. 2022;132(12).

37. O’Shaughnessy-Kirwan A, Signolet J, Costello I, Gharbi S, Hendrich B. Constraint of gene expression by the chromatin remodelling protein CHD4 facilitates lineage specification. Development. 2015;142(15):2586–97.

38. Wilczewski CM, Hepperla AJ, Shimbo T, Wasson L, Robbe ZL, Davis IJ, et al. CHD4 and the NuRD complex directly control cardiac sarcomere formation. Proc Natl Acad Sci U S A. 2018;115(26):6727–32.

39. Weiss K, Terhal PA, Cohen L, Bruccoleri M, Irving M, Martinez AF, et al. De Novo Mutations in CHD4, an ATP-Dependent Chromatin Remodeler Gene, Cause an Intellectual Disability Syndrome with Distinctive Dysmorphisms. Am J Hum Genet. 2016;99(4):934–41.

40. Weiss K, Lazar HP, Kurolap A, Martinez AF, Paperna T, Cohen L, et al. The CHD4-related syndrome: a comprehensive investigation of the clinical spectrum, genotype-phenotype correlations, and molecular basis. Genet Med. 2020;22(2):389–97.

41. Li Y, Liu Q, McGrail DJ, Dai H, Li K, Lin SY. CHD4 mutations promote endometrial cancer stemness by activating TGF-beta signaling. Am J Cancer Res. 2018;8(5):903–14.

42. Zhang Q, Zhu F, Tong Y, Shi D, Zhang J. CHD4 R975H mutant activates tumorigenic pathways and promotes stemness and M2-like macrophage polarization in endometrial cancer. Sci Rep. 2024;14(1):18617.

43. Daikoku T, Ogawa Y, Terakawa J, Ogawa A, DeFalco T, Dey SK. Lactoferrin-iCre: a new mouse line to study uterine epithelial gene function. Endocrinology. 2014;155(7):2718–24.

44. Hebeda CB, Savioli AC, Scharf P, de Paula-Silva M, Gil CD, Farsky SHP, et al. Neutrophil depletion in the pre-implantation phase impairs pregnancy index, placenta and fetus development. Front Immunol. 2022;13:969336.

45. Mills AA. The Chromodomain Helicase DNA-Binding Chromatin Remodelers: Family Traits that Protect from and Promote Cancer. Cold Spring Harb Perspect Med. 2017;7(4).

46. Hoffmeister H, Fuchs A, Erdel F, Pinz S, Grobner-Ferreira R, Bruckmann A, et al. CHD3 and CHD4 form distinct NuRD complexes with different yet overlapping functionality. Nucleic Acids Res. 2017;45(18):10534–54.

47. Nitarska J, Smith JG, Sherlock WT, Hillege MM, Nott A, Barshop WD, et al. A Functional Switch of NuRD Chromatin Remodeling Complex Subunits Regulates Mouse Cortical Development. Cell Rep. 2016;17(6):1683–98.

48. Kovac K, Sauer A, Macinkovic I, Awe S, Finkernagel F, Hoffmeister H, et al. Tumour-associated missense mutations in the dMi-2 ATPase alters nucleosome remodelling properties in a mutation-specific manner. Nat Commun. 2018;9(1):2112.

49. Doyle B, Morton JP, Delaney DW, Ridgway RA, Wilkins JA, Sansom OJ. p53 mutation and loss have different effects on tumourigenesis in a novel mouse model of pleomorphic rhabdomyosarcoma. J Pathol. 2010;222(2):129–37.

50. Guan B, Rahmanto YS, Wu RC, Wang Y, Wang Z, Wang TL, et al. Roles of deletion of Arid1a, a tumor suppressor, in mouse ovarian tumorigenesis. J Natl Cancer Inst. 2014;106(7).

51. Garcia-Sanz P, Trivino JC, Mota A, Perez Lopez M, Colas E, Rojo-Sebastian A, et al. Chromatin remodelling and DNA repair genes are frequently mutated in endometrioid endometrial carcinoma. Int J Cancer. 2017;140(7):1551–63.

